# chromMAGMA: regulatory element-centric interrogation of risk variants

**DOI:** 10.1101/2022.01.21.477270

**Authors:** Robbin Nameki, Anamay Shetty, Eileen Dareng, Jonathan Tyrer, Xianzhi Lin, the Ovarian Cancer Association Consortium, Paul Pharoah, Rosario I. Corona, Siddhartha Kar, Kate Lawrenson

## Abstract

Genome-wide association studies (GWASs) have identified thousands of genetic variants associated with common polygenic traits. The candidate causal risk variants reside almost exclusively in noncoding regions of the genome and the underlying mechanisms remain elusive for most. Innovative approaches are necessary to understand their biological function. Multimarker analysis of genomic annotation (MAGMA) is a widely used program that nominates candidate risk genes by mapping single-nucleotide polymorphism (SNP) summary statistics from genome-wide association studies to gene bodies. We augmented MAGMA into chromatin-MAGMA (chromMAGMA), a novel method to nominate candidate risk genes based on the presence of risk variants within noncoding regulatory elements (REs). We applied chromMAGMA to a genetic susceptibility dataset for epithelial ovarian cancer (EOC), a rare gynecologic malignancy characterized by high mortality. Disease-specific RE landscapes were defined using H3K27ac chromatin immunoprecipitation-sequence data. This identified 155 unique candidate EOC risk genes across five EOC histotypes; 83% (105/127) of high-grade serous ovarian cancer risk genes had not previously been implicated in this EOC histotype. Risk genes nominated by chromMAGMA converged on mRNA splicing and transcriptional dysregulation pathways. chromMAGMA is a pipeline that nominates candidate risk genes through a gene regulation-focused approach and helps interpret the biological mechanism of noncoding risk variants in complex diseases.

## INTRODUCTION

Multi-marker Analysis of Genomic Annotation, or MAGMA (de Leeuw et al. 2015) uses multiple regression to group raw or summary SNP statistics from GWASs to the level of genes while accounting for linkage disequilibrium (LD) between variants. Instead of testing millions of variants individually, MAGMA reduces the multiple testing burden by performing gene level analyses and has emerged as a powerful approach for the discovery of candidate genes and pathways associated with risk of complex traits (Wray et al. 2018; Demontis et al. 2019; Jansen et al. 2019). MAGMA captures SNPs positionally mapped to gene-bodies; however, many studies have now shown that noncoding tissue-specific REs (such as transcriptional enhancers marked by H3K27ac) are enriched for risk SNPs, and risk REs often interact with genes hundreds of kilobases away (Gerasimova et al. 2013, 2019; Jones et al. 2020; Nasser et al. 2021). Identifying candidate risk REs and the gene(s) they regulate remains a major bottleneck in the post-GWAS field. We therefore created a bioinformatic tool termed ‘chromatin-MAGMA’, or chromMAGMA, a pipeline that augments MAGMA to infer the target gene of noncoding risk variants based on user-inputted disease-relevant REs and RE-to-gene maps.

Here we tested chromMAGMA in epithelial ovarian cancer (EOC), a deadly disease with approximately 22,240 new cases and 14,070 annual deaths in the US (Torre et al. 2018). EOC can be stratified into five main histologic subtypes (histotypes) - high-grade serous (HGSOC), low-grade serous (LGSOC), endometrioid (EnOC), clear cell (CCOC), and mucinous ovarian cancer (MOC) (Soslow 2008; Torre et al. 2018). Each histotype is characterized by distinct molecular drivers, clinicopathologic features, and distinct germline genetic risk variants (Jones et al. 2017; Nameki et al. 2021). Of the 39 known unique EOC susceptibility loci (P-value < 5×10^-8^) identified through GWASs, 9 are associated with risk of HGSOC, 5 with risk of LGSOC, 4 with risk of MOC, and 1 with risk of EnOC. 20 loci are associated with all invasive disease, or a combination of one or more histotypes (Song et al. 2009; Bolton et al. 2010; Goode et al. 2010; Bojesen et al. 2013; Pharoah et al. 2013; Kelemen et al. 2015; Kuchenbaecker et al. 2015; Lawrenson et al. 2015; Kar et al. 2016; Lawrenson et al. 2016; Jones et al. 2017; Phelan et al. 2017). These genome-wide significant risk loci represent a fraction of all narrow-sense heritability in EOC and it is predicted that additional SNPs also contribute to disease susceptibility (Manolio et al. 2009; Boyle et al. 2017). Innovative approaches are needed to deconvolute additional true risk loci falling below genome-wide significance from false positives due to limited power, particularly for the rarer histotypes.

We applied chromMAGMA to EOC, inputting histotype-specific GWAS summary statistics, REs identified by H3K27ac chromatin immunoprecipitation-sequencing (ChIP-Seq) of Müllerian tissues, and RE-to-gene maps from the GeneHancer database (Corona et al. 2019). ChromMAGMA highlighted mRNA splicing and transcriptional dysregulation in EOC risk. In addition, lineage-specifying transcription factors (TFs) marked by super-enhancers (large stretches of H3K27ac) were particularly enriched for EOC risk associations based on chromMAGMA analyses and are likely to represent the nexus of noncoding EOC risk and transcriptional dysregulation. Overall chromMAGMA offers a flexible, gene regulation-focused approach to nominate noncoding regulatory elements and target genes involved in risk of polygenic traits.

## RESULTS

### chromMAGMA maps risk-associated, active regulatory elements to target genes

To identify candidate risk REs and associated genes for polygenic traits, we built the chromMAGMA pipeline by modifying the pre-processing and processing steps of conventional MAGMA and tested its performance using GWAS summary statistics and epigenome data for epithelial ovarian cancer (Supplementary Table 1). First, the genome is trimmed to only include regions annotated as high-confidence active REs from the GeneHancer database (Fishilevich et al. 2017). This reduces the genome from three billion base pairs (bp) to ~400 million bp (Figure 1a). Since GeneHancer includes data from 46 tissue types, for an EOC-specific analysis we further restricted the universe of GeneHancer REs to those regions marked by H3K27ac in normal and malignant Müllerian tissues and cell lines (Coetzee et al. 2015; Fishilevich et al. 2017; Corona et al. 2019). This created a final genome of ~200 million base pairs containing only regions of active chromatin identified in ovarian cancer-relevant tissues. GWAS SNP identifiers (reference SNP cluster identifiers, rsIDs) from 6 histotype-specific GWAS summary statistics (CCOC, EnOC, HGSOC, LGSOC, MOC, and NMOC – a dataset consisting of all samples except for MOC) (Coetzee et al. 2021) (Figure 1b) were then positionally mapped to the aforementioned RE dataset by applying the MAGMA annotation command (Methods). The SNP rsID-to-RE annotation was then processed for gene-level analysis using MAGMA (Methods) alongside EOC GWAS SNP summary statistics (P-values) and 1000 Genome European panel reference LD data (1000 Genomes Project Consortium et al. 2015). As multiple REs can regulate one gene (Peng and Zhang 2018), each gene was assigned the P-value of the most significant RE.

**Figure 1.**
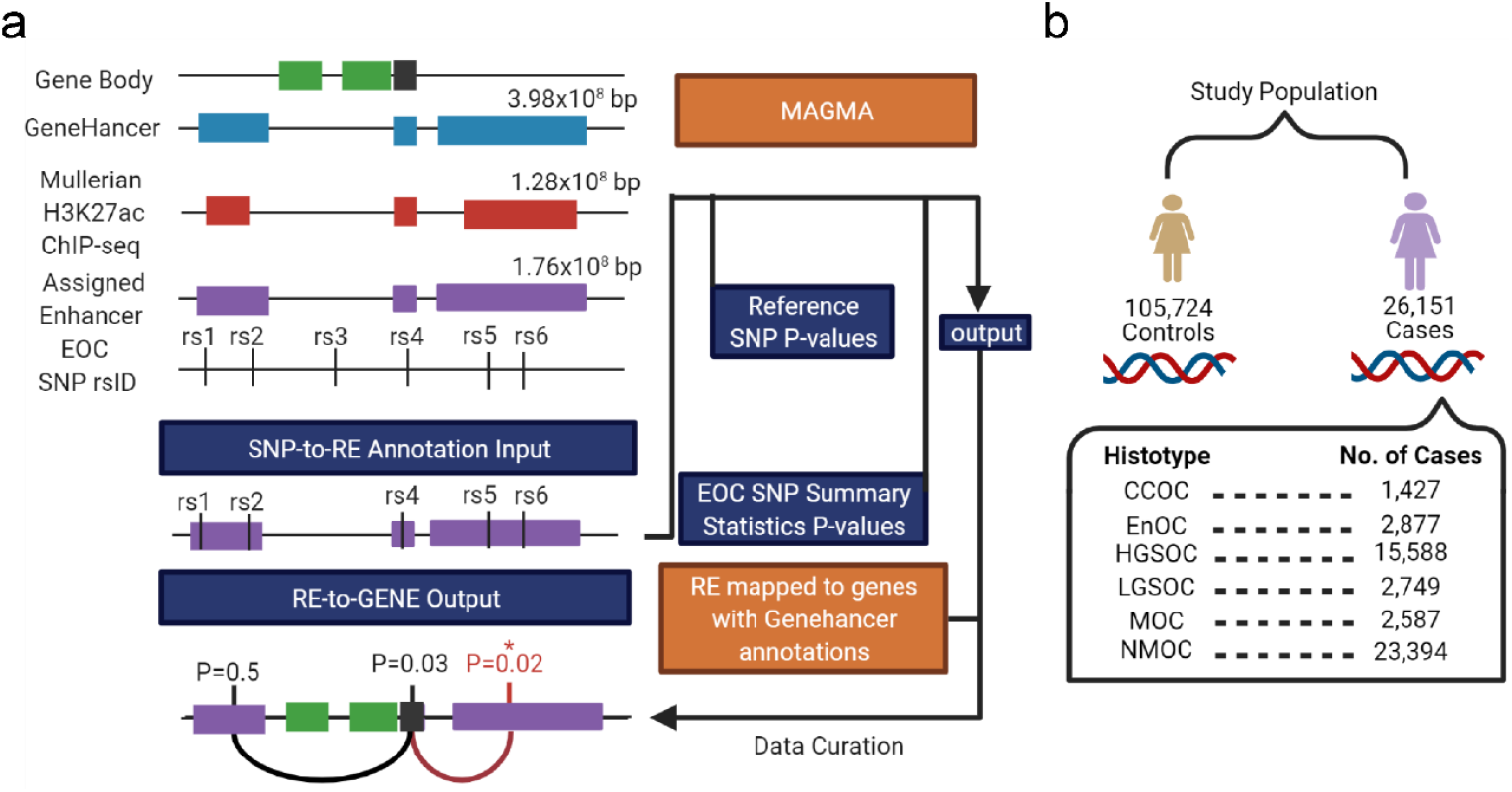
Applying chromMAGMA to EOC risk. **a)** Overview of the chromMAGMA approach. The GeneHancer database of regulatory elements (and linked genes) was limited to REs detected in Müllerian tissues. An EOC SNP rsID-to-RE-to-gene annotation list was created and used for gene-level analysis using the MAGMA model, along with EOC GWAS summary statistic P-values and reference linkage disequilibrium correlations from the European ancestry subset of the 1000 Genomes reference panel. Since multiple REs can be associated with one gene, the RE with the most significant P-value represents each gene. **b)** Study population of EOC GWAS dataset from Coetzee et al. 2021 CCOC, clear cell ovarian cancer; EnOC, endometrioid ovarian cancer; HGSOC, high-grade serous ovarian cancer; MOC, mucinous ovarian cancer; NMOC, all non-mucinous ovarian cancers (Coetzee et al. 2021).

Since REs marked by H3K27ac can identify both transcriptionally active promoters and enhancers, we stratified REs into promoters (defined as 1000bp upstream and 100bp downstream of a transcription start site) or candidate enhancers (all other regions of active chromatin) (Methods). This identified an average of 9,682 risk-associated REs for each histotype (range: 9,624-9,713) assigned to 17,435 protein-coding genes, of which 38% of the REs (3,703/9,682; range: 3,669-3,731) were active promoters and 62% (5,979/9,682; range: 5,966-6,002) were enhancers. The enhancer-to-promoter distance varied widely, with an average distance of 187,647 bp (range: 2 bp - 5 Mbp; SD +/- 230,272 bp) between enhancer start and transcription start sites (Supplementary Figure 1)

We ran conventional MAGMA alongside chromMAGMA to compare risk genes implicated by the two methods (Supplementary Table 2). In MAGMA and chromMAGMA, the P-value is calculated in a two-step process: first the SNP matrix is projected into a smaller set of principal components to remove the effects of highly correlated SNPs; second these principal components are used in a linear regression whose outputs (feature-wise enrichment for significant SNPs) are tested for statistical significance using an F-test (de Leeuw et al. 2015). Considering all protein coding genes, the average chromMAGMA P-value was significantly lower compared to MAGMA across EOC histotypes (P-value < 0.001 for all histotypes; Welch two-sample T-test), consistent with previous evidence that REs, but not protein coding exons, are enriched for risk-associated SNPs (2019). After Bonferroni correction to account for the total number of genes tested in each histotype-specific analysis, we identified 68 unique significant genes in MAGMA and 155 unique significant genes in chromMAGMA, with 56 genes identified by both methods (MAGMA Bonferroni corrected P-value < 2.70×10^-6^; chromMAGMA Bonferroni corrected P-value < 2.87×10^-6^). The number of genes identified by histotype ranged from 0 (EnOC) to 53 (all non-mucinous cancers, NMOC) significant genes in MAGMA and 0 (EnOC) to 131 (NMOC) significant genes using chromMAGMA (Figure 2a). Disparity in the number of significant genes by histotype is likely due to power, as HGSOC and NMOC represents a majority of the overall sample size in the EOC GWAS.

**Figure 2.**
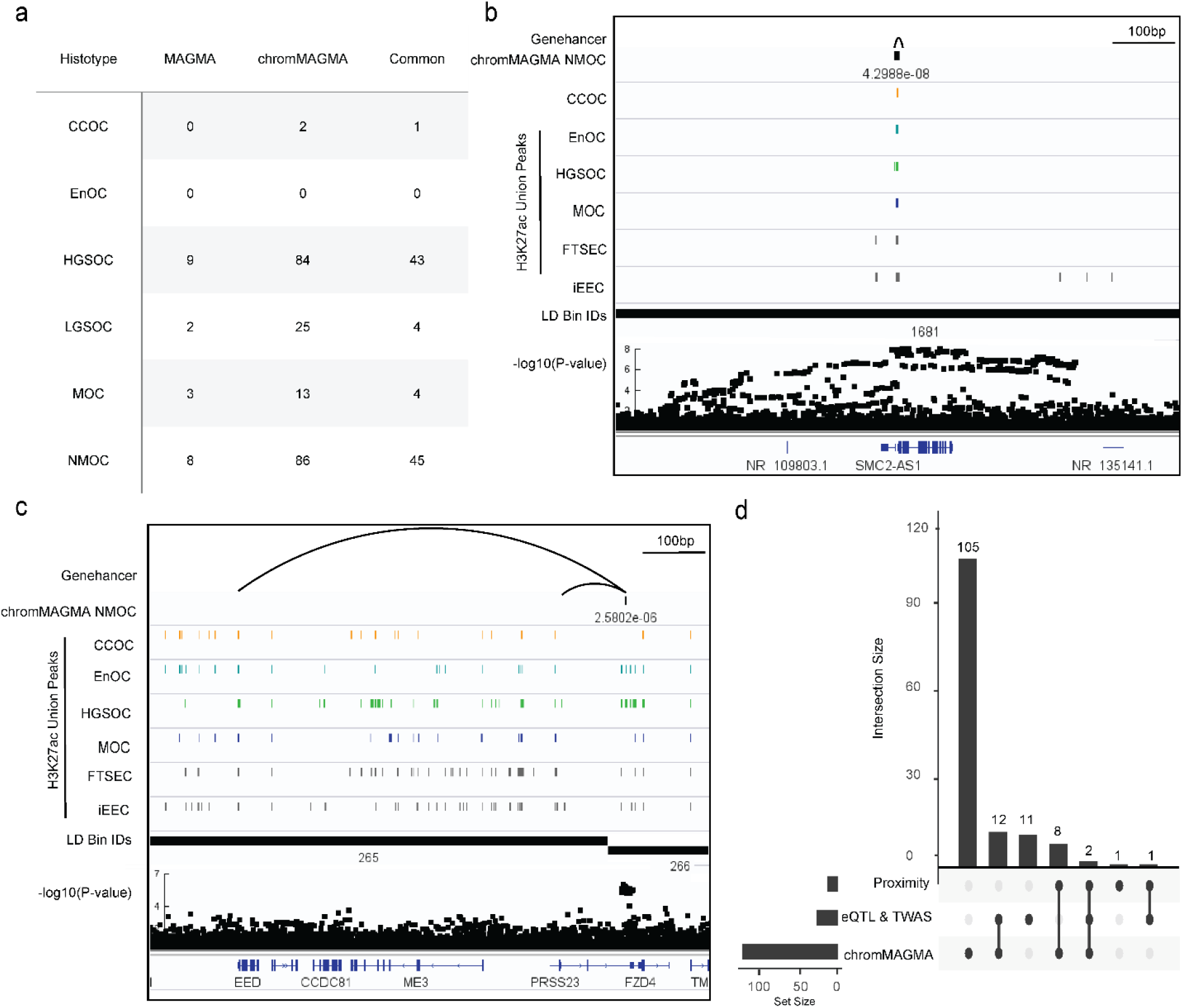
ChromMAGMA identifies risk genes for epithelial ovarian cancer through risk SNPs coinciding with regulatory elements. **a)** Candidate risk genes identified by conventional MAGMA and chromMAGMA across EOC histotypes (adjusted P-value < 0.05). **b)** Locus view displaying the NMOC chromMAGMA RE-to-gene association for *SMC2*. LD boundaries and GWAS SNP associations (-log_10_(P-value)) are shown. **c)** Locus view displaying the NMOC chromMAGMA RE-to-gene association for *PRSS23* and *EED*. **d)** UpSet plot of chromMAGMA genome-wide significant genes in HGSOC and alternate approaches to nominate candidate risk genes. Proximity, lead variants labeled as genome-wide significant (P < 5×10^-8^) assigned to genes based on nearest transcription start site; eQTL, *cis*-expression quantitative trait loci; TWAS, transcriptome-wide association studies; CCOC, clear cell ovarian cancer; EnOC, endometrioid ovarian cancer; HGSOC, high-grade serous ovarian cancer; LGSOC, low-grade serous ovarian cancer; MOC, mucinous ovarian cancer; NMOC, all non-mucinous ovarian cancers.

Gene-dense GWAS loci at genome-wide significance account for many of the risk genes identified by MAGMA and chromMAGMA (Berisa and Pickrell 2016). For example, genes on chromosome 17 are overrepresented in MAGMA and chromMAGMA in EOC (26/68 and 52/155 unique genes respectively) likely due to the presence of two genome-wide significant loci in this chromosome and the high degree of LD due to an inversion at 17q31 (Jones et al. 2017).We divided the genome into distinct bins based on LD to identify instances where chromMAGMA nominates candidate risk REs within novel LD bins, scenarios where the same candidate gene could not readily be identified through MAGMA. Using this approach, twenty-nine unique genes were identified as candidate risk genes only in chromMAGMA (Supplementary Table 3). Using chromMAGMA NMOC as an example, a significant promoter (P-value 4.3×10^-8^) at a known breast and ovarian cancer genome-wide significant risk locus at chromosome 9q31 is assigned to *SMC2*; whereas in MAGMA, *SMC2* is not significant (P-value 1.9×10^-3^) (Kar et al. 2016) (Figure 2b). Other candidates not previously implicated in EOC risk such as *PRSS23* (P-value 2.6×10^-6^), a serine-protease regulated by HGSOC biomarker PAX8 (Adler et al. 2017) were also identified (Figure 2c). The same RE assigned to *PRSS23* interacts across an LD boundary with the promoter of *EED*. EED is a component of the polycomb repressor complex involved in the pathogenesis of numerous cancer types (Kim et al. 2013).

We next compared chromMAGMA risk genes with candidate susceptibility genes nominated by alternative approaches. This analysis was limited to HGSOC as it is the most common and well-studied EOC subtype. Chromosome conformation capture assays have identified candidate susceptibility genes previously undiscovered based on proximity to the nearest gene promoter. So far three GWAS significant loci at 11q31, 8q24, and 19p13 originally mapped by proximity to *HOXD3, PVT1*, and *BABAM1* was found, *via* chromosome conformation capture assays to interact with *HOXD9, MYC*, and *ABHD8* respectively (Grisanzio and Freedman 2010; Lawrenson et al. 2015, 2016). chromMAGMA nominated all three as candidate susceptibility genes in HGSOC (*HOXD9*, P-value 1.52×10^-12^; *MYC*, P-value 6.39×10^-11^ and *ABHD8*, P-value 3.9×10^-16^). As chromMAGMA also identifies risk variants that may impact short-range enhancer-promoter interactions, promoters, and intronic enhancers, we reasoned that it should also be able to capture susceptibility genes identified through traditional GWAS based on closest proximity. Indeed, 10 out of 12 (83 %) genes previously labeled as genome-wide significant based on proximity to a lead variant overlapped with chromMAGMA nominated genes (P-value < 5×10^-8^) (Figure 2d). *Cis*-expression quantitative trait loci (eQTL) and transcriptome-wide association studies (TWASs) integrate genotype data with gene expression to identify candidate genes associated with disease risk. To date, 26 candidate genes have been identified as HGSOC candidate risk genes using these methods (Lawrenson et al. 2015; Lu et al. 2018; Kar et al. 2020); 16 out of 26 genes (62%) previously identified by HGSOC eQTL or TWAS analyses were also nominated by chromMAGMA (Figure 2d). chromMAGMA identified 105 additional genes previously not implicated in HGSOC risk. 22/105 of these genes had long range interactions (> 500 kilobases) with the risk RE, highlighting how chromMAGMA can identify candidate genes transcriptionally impacted by noncoding risk SNPs that are hundreds of kilobases away. For example, the longest promoter-risk RE interaction identified in this analysis was between the *GMPS* gene (P-value 5×10^-10^) and an associated RE 9.4 kilobases away in linear genomic distance. Overall, chromMAGMA nominates candidate risk genes that are consistent with alternate methods, but also implicates additional genes in HGSOC susceptibility through risk SNPs with their upstream active regulatory elements.

### chromMAGMA implicates splicing and gene regulation in EOC risk

To identify pathways regulated by risk-associated regulatory elements, we conducted gene set enrichment analysis to ask whether any gene sets from the Gene Ontology (GO) database were enriched in ranked gene-level associations (based on descending order of -log_10_(P-value)) from MAGMA and chromMAGMA. This approach allows for the investigation of sets of genes without the need to assign arbitrary P-value cutoffs. Since one RE can be assigned to multiple genes in the chromMAGMA gene-level association, gene ranks were weighted using a mRNA expression dataset comprised of disease-relevant primary tissue samples (Corona et al. 2019) to generate a ranked list in which highly expressed genes are ranked higher than relatively lower expressed genes associated with the same RE (Methods). Pathway enrichment analysis with the chromMAGMA-derived gene list identified 140 common pathways across all histotypes, of which 7 were related to mRNA splicing and processing (considering only pathways with positive normalized enrichment scores and adjusted P-value < 0.05) (Figure 3a, Supplementary Table 4). Spliceosome factors *CHERP* and *EFTUD2* were the top 2 (out of 349) most significant genes related to mRNA splicing in the weighted chromMAGMA gene list. In addition, 20 of the common pathways were terms related to transcription or chromatin, including RNA polymerase II activity and transcription factor activities (Figure 3a). DNA-binding Transcription Factors *SIN3B* and *NFE2*L1 were the top 2 most significant genes (out of 1746) in the weighted chromMAGMA gene list and the transcription mediator complex coactivator *MED26* ranked 3^rd^. Histotype-specific pathways were also observed for CCOC (195 pathways), EnOC (58 pathways), HGSOC (17 pathways), LGSOC (30 pathways) and MOC (70 pathways) (Figure 3b). In contrast, pathway gene set enrichment with conventional MAGMA had no enriched pathways that passed the P <0.05 (after adjustment for multiple comparisons) threshold across all histotypes.

**Figure 3.**
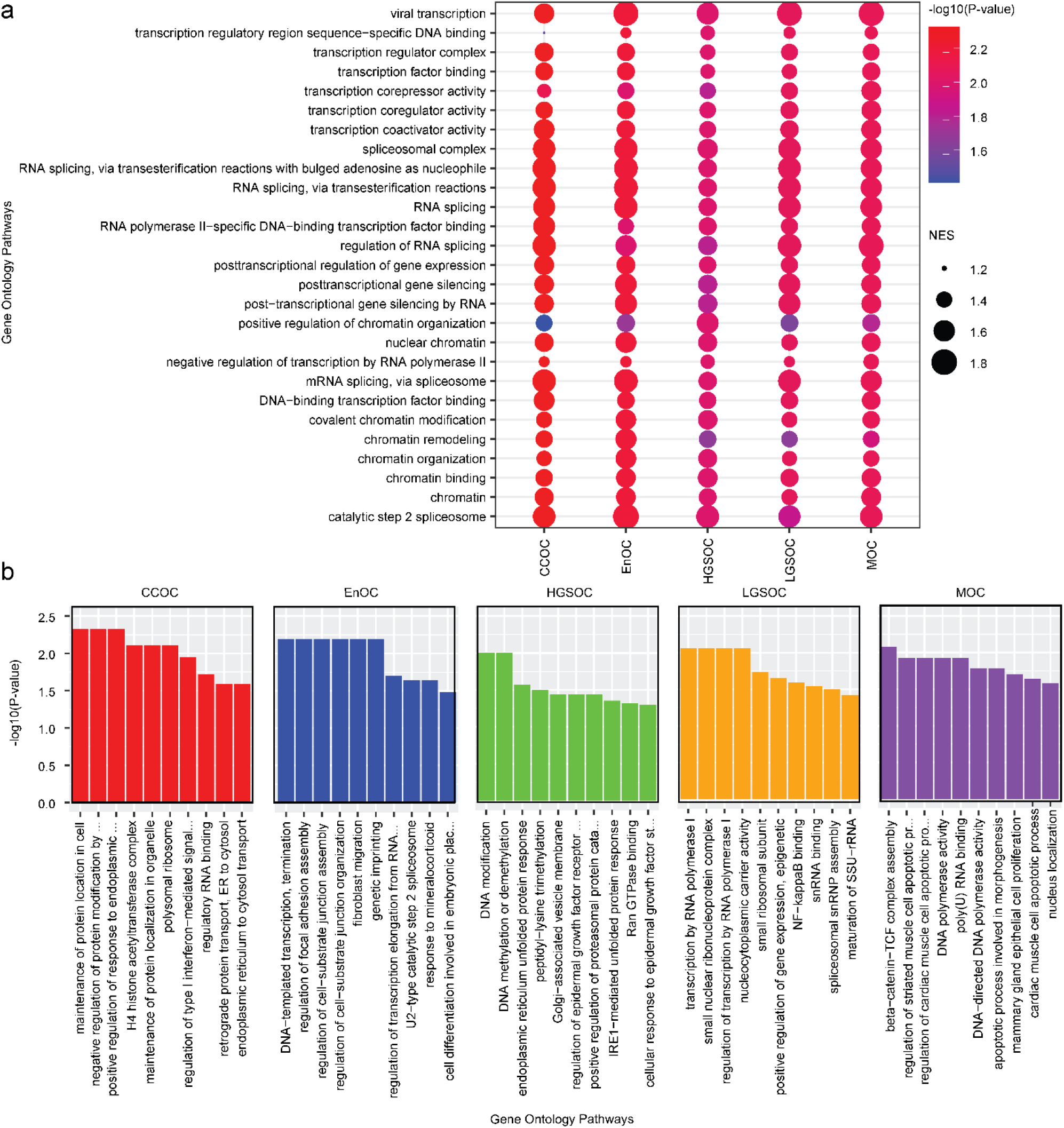
chromMAGMA identifies histotype-specific as well as common pathways involved in EOC risk. **a)** Dot plot representing transcription, splicing, and chromatin related pathways that were enriched in risk genes nominated in all histotypes by chromMAGMA. **b)** Bar plots representing the top 10 histotype-specific chromMAGMA pathways based on normalized enrichment score. NES, normalized enrichment score; CCOC, clear cell ovarian cancer; EnOC, endometrioid ovarian cancer; HGSOC, high-grade serous ovarian cancer; LGSOC, low-grade serous ovarian cancer; MOC, mucinous ovarian cancer

### Super-enhancer-associated transcription factors are associated with EOC risk

Cancer cells are often dependent on lineage-specifying TFs whose expression is propelled by large clusters of enhancers termed super-enhancers or stretch enhancers (Bradner et al. 2017). Of 1,671 known human TFs, 257, 220, 202, and 247 TFs are associated with super-enhancers in CCOC, EnOC, HGSOC, and MOC respectively (Supplementary Table 5). Using chromMAGMA we identified super-enhancer-associated TFs as enriched for association with risk at P-value < 0.05; FDR (q-value) < 0.25 for all histotypes except LGSOC, as LGSOC tissue H3K27ac ChIP-seq data were not available (Figure 4a) (Hnisz et al. 2013; Whyte et al. 2013; Corona et al. 2019). By contrast, super-enhancer associated TFs were only significantly enriched for HGSOC risk (P-value < 0.05; FDR q-value < 0.25) when using gene-level statistics derived from conventional MAGMA. Leading-edge analysis was performed to identify the super-enhancer associated TFs overrepresented in the top ranks of chromMAGMA gene-level associations. TFs previously implicated in epithelial ovarian cancer development including 6/14 candidate master regulators for HGSOC based on a recent pan-cancer gene expression analysis were implicated in EOC risk (Supplementary Table 6). Three of these factors (PAX8, SOX17, and MECOM) are functionally validated master regulators of HGSOC development (Figure 4b) (Reddy et al. 2019). *HNF1B*, a CCOC biomarker and a key regulator of CCOC tumorigenesis (Cuff et al. 2013; Li et al. 2015), was also on the leading edge of the clear cell ovarian cancer analysis (Figure 4b).

**Figure 4.**
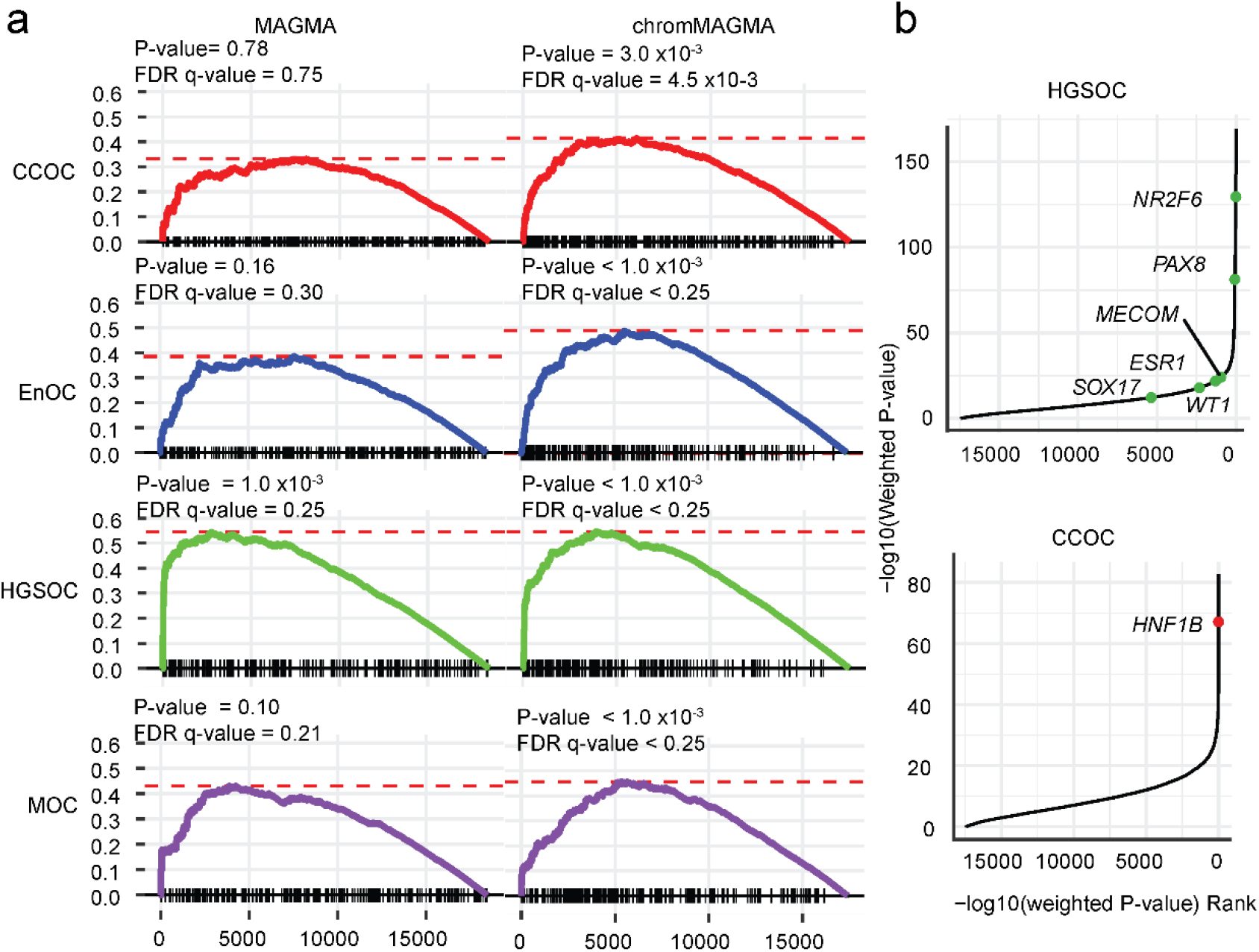
Super-enhancers upstream of transcription factors are associated with histotype-specific EOC risk. a) Gene-set enrichment plot of super-enhancer associated TFs from each EOC histotype from Corona et al. 2020 in conventional MAGMA and chromMAGMA. **b)** Gene -log_10_(P-value) *versus* gene rank based on - log10(P-value) with known genes implicated in CCOC and HGSOC from the leading-edge list of the superenhancer associated TF gene-set enrichment analysis highlighted. CCOC, clear cell ovarian cancer; EnOC, endometrioid ovarian cancer; HGSOC, high-grade serous ovarian cancer; LGSOC, low-grade serous ovarian cancer; MOC, mucinous ovarian cancer; NMOC, all non-mucinous ovarian cancers.

### Gene set enrichment analysis of TF cistromes

In addition to TFs being the target of risk SNPs, noncoding SNPs may also impact disease risk by modifying TF binding within enhancers to impact gene expression (Oldridge et al. 2015; Kandaswamy et al. 2016; Huo et al. 2019). Therefore, we asked whether target genes of specific TFs are disproportionally impacted by EOC risk SNPs in chromMAGMA. For this analysis we asked if TF-specific gene sets in the Molecular Signatures database (MsigDB) are enriched in the ranked gene list from chromMAGMA (Subramanian et al. 2005; Liberzon et al. 2011) (Supplementary Table 7). TF targets are defined as genes with motifs located within 4 kb around their transcription start sites by MsigDB.

We first explored the PAX8 target gene sets in HGSOC, as we have previously identified PAX8 target gene sets to be enriched in this histotype(Kar et al. 2017). PAX8 is represented by two gene sets in MsigDB - PAX8_B contains 106 genes, PAX8_01 contains 39 genes, with 23 genes in common across the two sets. When ranked by the normalized enrichment score, PAX8_B ranked 24/573 (P-value =0.032, FDR q-value = 0.098) and PAX8_01 ranked 42/573 (P-value =0.13, FDR q-value = 0.34) in MAGMA. With chromMAGMA, the PAX8 target gene sets ranked higher, with PAX8_B ranked 1/573 (P-value < 1.0 x10^-3^, FDR q-value = 0.115), and PAX8_01 ranked 12/573 (P-value =0.040, FDR q-value = 0.18). We also explored chromMAGMA performed for NMOC, and this analysis identified targets of EVI1, also known as MDS1 and EVI1 Complex Locus (MECOM) as a significant gene set not identified in MAGMA (MsigDB-EVI1_05 P-value = 1.0 x10^-3^, FDR q-value = 0.17; MsigDB-EVI1_04; P-value = .041, FDR q-value = 0.15; MsigDB-EVI1_03; P-value= 4.6 x10-2, FDR q-value = 0.229). MECOM is a known master regulator TF in HGSOC that is functionally involved in disease pathogenesis (Reddy et al. 2019; Bleu et al. 2021). Leading-edge analysis was performed for MsigDB-PAX8_B and MsigDB-EVI1_05 to identify the candidate susceptibility genes potentially regulated by PAX8 and MECOM. This analysis identified 29 and 49 candidate susceptibility genes regulated by PAX8 and MECOM, respectively. Interestingly, *HOXB5, HOXB7, HOXB8*, and *NEUROD6* genes were common target genes between the two factors. *HOXB5, HOXB7*, and *HOXB8* are homeobox superfamily TFs that are particularly highly expressed in HGSOC and associated with poor survival (Idaikkadar et al. 2019).

We then used chromMAGMA to discover additional TFs not previously implicated in EOC risk with histotype specificity in consideration. Since one transcription factor can be represented by multiple gene sets, the gene set with the most significant P-value was chosen to represent each transcription factor. chromMAGMA performed for NMOC were excluded from this as it is a dataset consisting of all samples except for MOC. Considering P-value <0.05 & FDR cutoff of < 0.25, we identified 113 transcription factors implicated in EOC risk. Of these 113 transcription factors, 13 were specific to CCOC, 4 to EnOC, 5 to HGSOC, 7 to LGSOC, 9 to MOC and 7 common across all 5 histotypes (Figure 5a, Supplementary Figure 5a). *SOX9* was identified as a CCOC specific TF in which its downstream regulatory targets are enriched for risk SNPs. A recent single-cell RNA sequencing study of the human endometrium (hypothesized tissue-of-origin for CCOC) grouped SOX9 positive epithelial cells of the endometrium as a regenerating and proliferative subset (Garcia-Alonso et al. 2021).

**Figure 5.**
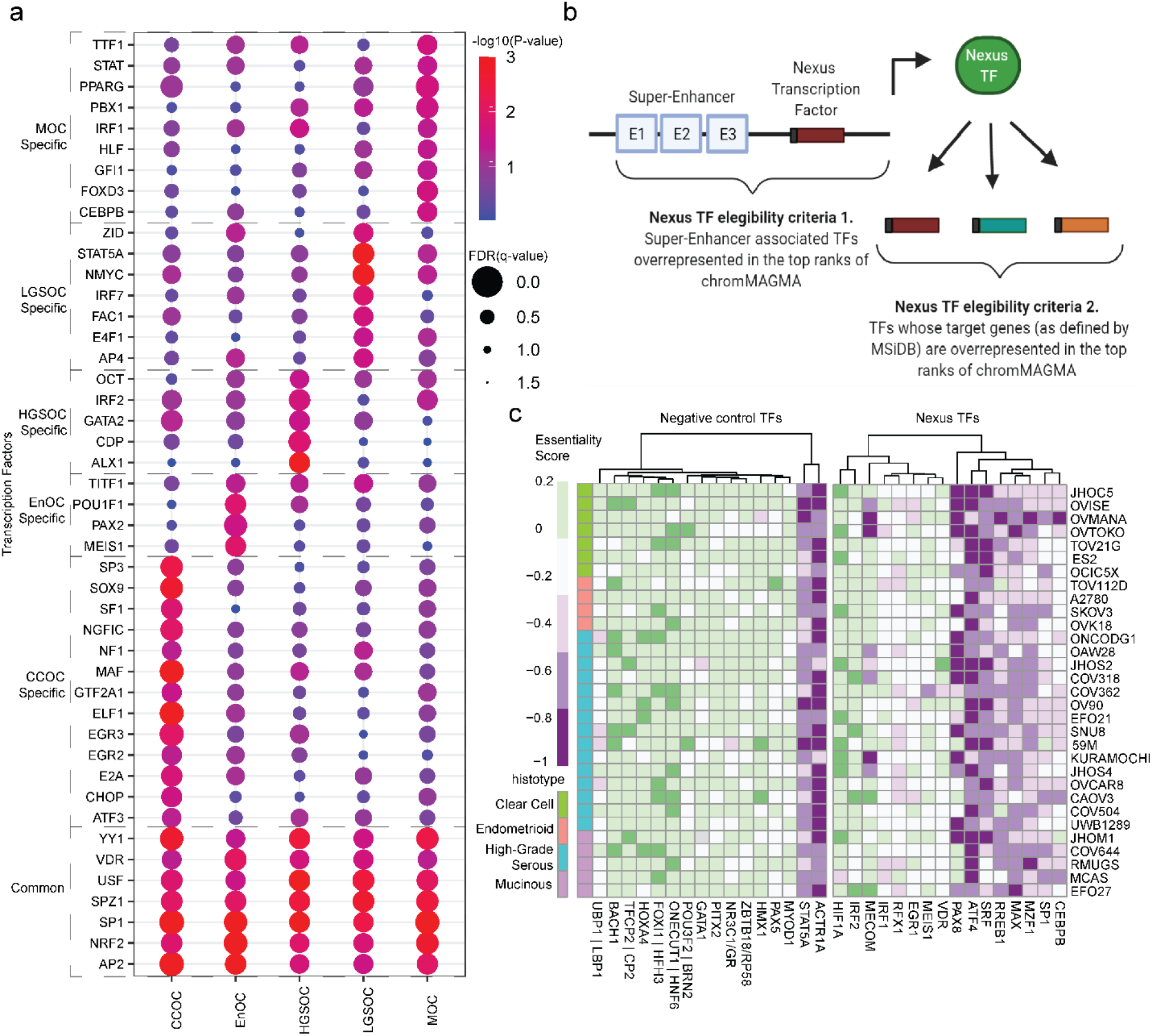
Transcription factor networks in EOC risk. a) Genome-wide significant Molecular Signatures Databse transcription factor targets (MsigDB ‘TFT_legacy’) gene-sets (FDR < 0.05) compared between MAGMA and chromMAGMA across histotypes b) Schematic depiction of the definition of a ‘Nexus TF’. Top ranks = leading edge of the gene-set enrichment analysis. c) Heatmap displaying the essentiality score of Nexus TFs in EOC lines (data from Depmap.org). Columns clustered using unsupervised hierarchical clustering (method = X). CCOC, clear cell ovarian cancer; EnOC, endometrioid ovarian cancer; HGSOC, high-grade serous ovarian cancer; LGSOC, low-grade serous ovarian cancer; MOC, mucinous ovarian cancer; NMOC, all non-mucinous ovarian cancers.

Finally, we set to identify TFs that are likely to be directly regulated by risk SNPs and where risk SNPs also modify TF downstream binding, hereinafter termed as ‘nexus TFs’. Nexus TFs were defined as TFs that were (1) on the leading edge of the super-enhancer associated TF gene set enrichment analysis and (2) TF target gene sets from MsigDB that were significantly enriched in chromMAGMA for each respective histotype (Figure 5b, Supplementary Table 7). 16 TFs such as PAX8 were identified for HGSOC and EnOC, along with novel TFs implicated in EOC such as SP1, a TF implicated in a variety of biological processes across multiple cancer types (Vizcaíno et al. 2015). To explore the functional role of Nexus TFs, we leveraged a publicly available CRISPR-Cas9 knock-out screen that includes seven CCOC cell lines, four EnOC cell lines, 15 HGSOC cell lines, and five MOC cell line models (Meyers et al. 2017). Although there is heterogeneity across cell lines and histotypes, EOC lines are largely dependent on *RREB1, ATF4, MAX, PAX8, MZF1, SRF* (average essentiality scores ≤ −0.4; average score for pan-essential genes = −1) and *SP1, MECOM*, and *CEBPB* (average essentiality scores ≤ −0.3) (Figure 5c, Table 1). By comparison, negative control TFs that were (1) not on the leading edge of the super-enhancer associated TF gene set enrichment analysis and (2) bottom 16 of the TF target gene sets from MsigDB were less likely to be essential in EOC cell lines (Figure 5c). In total 9/16 nexus TFs showed at least modest dependency (average essentiality scores ≤ −0.3) in at least one histotype, compared to 2/16 negative control TFs. These results imply that TFs on the nexus of risk through genetic variation both in upstream REs and downstream binding sites can be identified in chromMAGMA and are often essential genes in EOC.

**Table 1.**
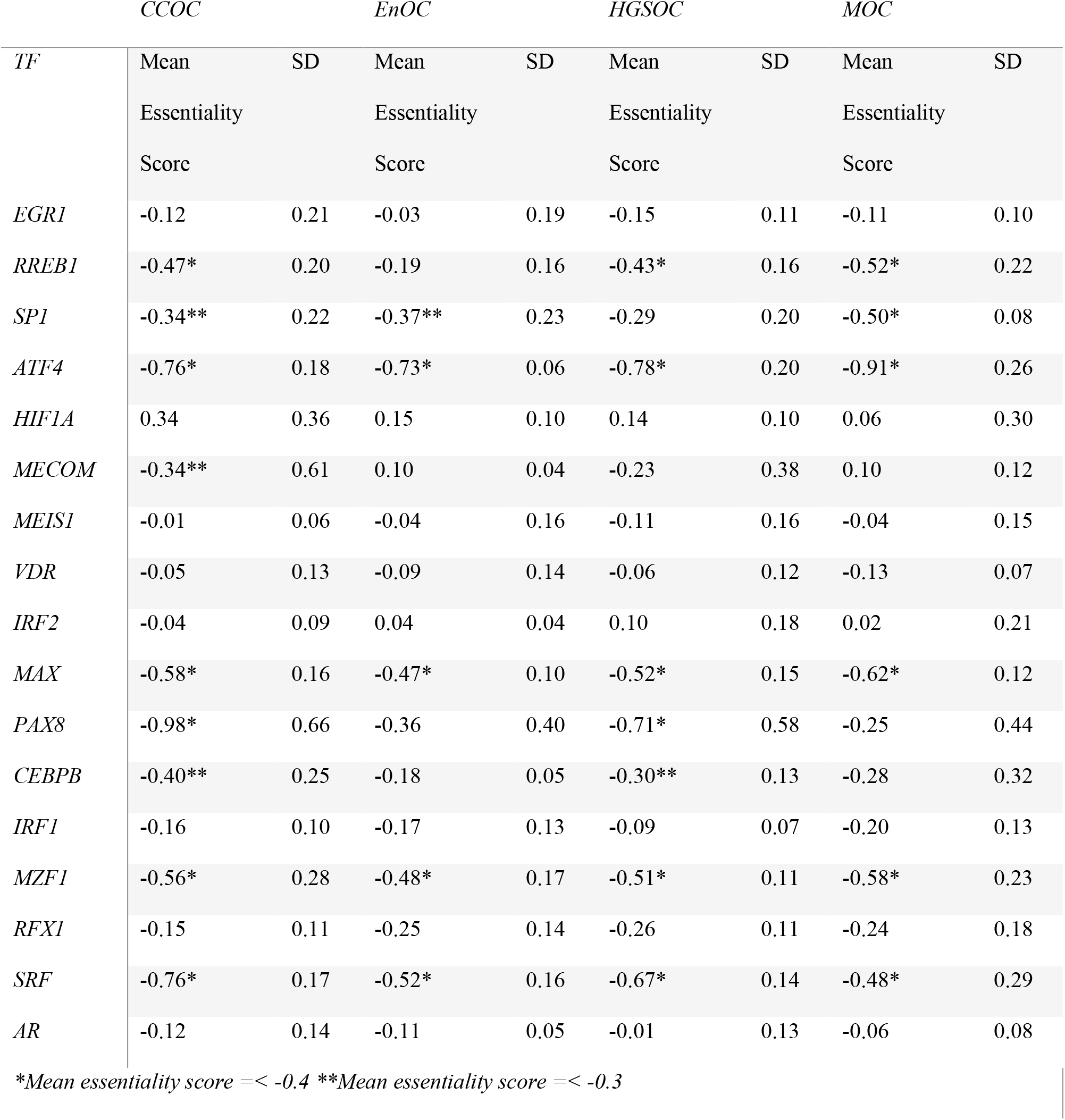
Average essentiality scores for Nexus TFs across EOC cell lines. Mean essentiality score represents the average essentiality score for all cell lines associated with each EOC histotype. CCOC, clear cell ovarian cancer; EnOC, endometrioid ovarian cancer; HGSOC, high-grade serous ovarian cancer; LGSOC, low-grade serous ovarian cancer; MOC, mucinous ovarian cancer; NMOC, all non-mucinous ovarian cancers; SD, standard deviation.

## DISCUSSION

Most common risk polymorphisms associated with complex traits in GWAS are located in the noncoding portion of the genome (Zhang and Lupski 2015). These noncoding risk polymorphisms likely modify the activity of noncoding regulatory elements to impact the expression of a target gene (or genes) that play a role in disease susceptibility (2019). Identifying the risk RE and target gene remain two main challenges in post-GWAS functional work, since REs outside of promoters are tissue-specific and can interact with transcription start sites over large linear genomic distances (Sanyal et al. 2012). Here we built chromatin-MAGMA, or ‘chromMAGMA’, to prioritize candidate risk REs and target genes based on the landscape of gene regulation in a specific tissue type. chromMAGMA first maps SNPs to user-defined tissue-specific regulatory element landscapes. REs are linked to putative target genes using the GeneHancer database, or any other resource. The collection of annotated active REs can then be interrogated at the gene level, or gene set level. Here we focused on epithelial data sets representing the tumor type of interest, plus likely precursor cell types; however analogous data sets for other cell types could also be interrogated, where they exist.

We tested the performance of chromMAGMA using the largest EOC GWAS dataset to date, consisting of 26,151 EOC cases and 105,724 controls (Coetzee et al. 2021) combined with disease-relevant Müllerian active REs and RE-to-gene contact maps from the GeneHancer database. We contrasted the RE-centric chromMAGMA to genes nominated by conventional MAGMA. Overall, chromMAGMA assigned lower P-values to genes compared to MAGMA, in line with evidence that SNPs are enriched in active REs, validating the overall premise of this approach (Gerasimova et al. 2013, 2019; Jones et al. 2020; Nasser et al. 2021). Orthogonal evidence to validate the chromMAGMA approach came from concordant results obtained when using chromMAGMA and alternative approaches to nominate candidate EOC susceptibility genes (including proximity, chromosome conformation capture assays, and quantitative trait locus-based analyses). Of particular note is that chromMAGMA identified previously validated candidate genes in scenarios where large genomic distances or multiple genes lie between the candidate causal risk SNPs and the risk gene. This highlights how the chromMAGMA approach represents an efficient route to candidate gene nomination that incorporates the benefits of popular existing methods, while avoiding some of the limitations associated with those techniques. For example, chromMAGMA circumvents the distance bias of both eQTL analyses (which are often only powered to identify local *cis* interactions) or analyses that leverage chromatin interactome data (which conversely cannot resolve short-range interactions, which poses a particular challenge in gene-dense regions).

In addition to validating known risk genes, chromMAGMA provided insights into EOC risk that have not been achieved using previous methods. Pathway analysis of the chromMAGMA ranked gene list revealed enrichments of mRNA processing and splicing pathways across all histotypes, indicating that noncoding risk SNPs falling on REs regulate genes within these pathways. While splicing events have been recently associated with EOC risk(Gusev et al. 2019), components of splicing machinery have not been implicated in EOC risk previously. Transcriptional regulation pathways were also enriched in risk genes highly ranked by chromMAGMA, particularly super-enhancer associated lineage specific factors (such as *PAX8* in HGSOC and *HNF1B* in CCOC). A study incorporating long-range, noncoding chromatin interactions from Hi-C with MAGMA (H-MAGMA) in 9 neuropsychiatric disorders also found common pathways in transcriptional regulation/RNA splicing(Sey et al. 2020). These results suggest that risk variants impacting such pathways may be common occurrences across complex traits. As TFs can be both targets and mediators of risk SNPs, we identified a set of ‘Nexus TFs’, i.e., transcription factors with oncogenic transcriptional properties that are enriched for risk variation both in its upstream *cis* regulatory element and in its downstream target binding sites. Gene dependency data prioritized nine transcription factors, which included master regulators of HGSOC - *PAX8* and *MECOM*. PAX8 and MECOM are known to co-occupy a majority of H3K27ac active regions in HGSOC, and may be contributing to the differential regulation of HGSOC-relevant risk genes (Reddy et al. 2019).

While our study used H3K27ac chromatin immunoprecipitation data - a widely available mark of active chromatin, other technologies and epigenetic marks - such as other histone post-translational modifications, transcription factor binding sites, open chromatin regions, and methylation profiles - are all compatible with chromMAGMA. In this study we used GeneHancer active regulatory-element-to-gene contact map data (Fishilevich et al. 2017). GeneHancer is the most comprehensive catalogue of gene-regulatory element associations currently available and is comprised of RE-to-gene maps represented by 46 tissue types. One limitation to this approach is that Müllerian tissues are not well represented in the GeneHancer database and could be missing interactions unique to gynecologic tissues. Alternative data types, such as *in silico* maps of RE-promoter interactions inferred from ATAC-seq data (Corces et al. 2018) or genome-wide data from epigenome and genome editing screens could be incorporated to create tissue-specific maps of gene-RE assignments. Another limitation of chromMAGMA is the necessary step of assigning a representative RE to a single gene for the generation of gene-level statistics. In this study, genes were mapped 1:1 to the RE with the most significant P-value. This step simplifies the biological complexity of multiple REs influencing a gene in an additive (Hay et al. 2016; Kawakami et al. 2021), or sometimes hierarchal manner (Shin et al. 2016; Huang et al. 2018), but for some genes, may miss a critical aspect of transcriptional regulation relevant to risk. Integration of chromMAGMA with data from perturbation (Fulco et al. 2019) and massively parallel reporter (Muerdter et al. 2015) assays may be a superior way to prioritize REs associated with each gene. Overall, chromMAGMA is a flexible approach that can be readily adapted to prioritize candidate risk genes and regulatory elements for a wide array of phenotypes.

## COMPUTATIONAL METHODS

### MAGMA

MAGMA uses the P-values of SNPs and local linkage disequilibrium to assign SNPs to gene locations, and then aggregates SNPs within the same gene body using a hypergeometric distribution (de Leeuw et al. 2015). These genes are then ranked by ranking the -log_10_(P-value). The greater the -log_10_(P-value), the greater the number and/or significance of GWAS SNPs lying within the interval of the gene. MAGMA requires two external data sources: a list of GWAS SNPs with associated P-values from that of GWAS, number of participants, and a list of annotations linking gene names to intervals in the genome.

GWAS data came from the OCAC consortium study of 26,151 cases and 105,274 controls participants (Coetzee et al. 2021). The GWAS data contained SNP P-values for five histotypes of ovarian cancer - high-grade serous, low-grade serous, clear cell, endometrioid, mucinous, and a composite category of all non-mucinous histotypes. Gene locations from the NCBI build 37. Significant genes were identified by filtering genes whose P-values were less than the Bonferroni corrected value of 2.70×10^-6^.

### chromMAGMA

A list of all REs (hg19) and corresponding gene targets was obtained from Genehancer (v4.7) a publicly available database of RE-to-gene maps (Fishilevich et al. 2017). Genehancer captures a broad universe of RE activity which we wished to reduce to those specific to ovarian cancer and precursor cell states. We used a dataset of H3K27ac peaks derived from clear cell (number of non-unique peaks = 119,549 peaks), endometrioid (125,743 peaks), high-grade serous (122,734 peaks), mucinous (131,655 peaks) ovarian tumor tissues, and samples from endometriosis epithelial (44,083 peaks), and normal fallopian tube secretory epithelial cell lines (43,734 peaks) (Coetzee et al. 2015; Corona et al. 2020). This was converted to hg19 using UCSC liftOver, duplicates were removed and the remainder were merged into 80,271 distinct intervals using bedtools v2.25.0 (Quinlan and Hall 2010). We then selected REs from Genehancers which overlapped with our H3K27ac intervals by at least one base pair.

We used the REs as the interval input into MAGMA to replace the gene intervals used by MAGMA. This generated a list of REs and their statistics. This list was then linked to the genes, where each gene was assigned the greatest -log_10_(P-value) from its REs. REs were defined as promoters based on the txdb.hsapiens.ucsc.hg19.knowngene database, and all non-promoters were labeled as candidate transcriptionally active enhancers. Significant genes were identified by filtering genes whose P-values were less than the Bonferroni corrected value of 2.87×10^-6^.

### Identifying proximal genes to GWAS genome-wide significant loci

All lead variants labeled as genome-wide significant (P < 5×10^-8^) in ovarian cancer by Jones et al. 2017 (Jones et al. 2017) were assigned to a gene based on nearest transcription start site.

### Generation of the gene list

Gene identifiers in chromMAGMA and MAGMA were curated by restricting to those identifiable as ‘ensembl_gene_id’, ‘external_gene_name’,’external_synonym’,’hgnc_symbol’, ‘entrez_gene_id’, and ‘uniprot_gn_symbol’ and filtered for genes labeled as ‘protein_coding’ from the BioMart portal (Smedley et al. 2015). For MAGMA, the maximal -log_10_(P-value) was then assigned to a gene, and simply ranked with -log_10_(P-value) in descending order. For chromMAGMA, ties in the -log_10_(P-value) were broken using the average expression of variance stabilization normalized primary CCOC, EnOC, HGSOC, MOC and fallopian tube secretory epithelium (Average Müllerian mRNA Expression) as described from Corona et al. 2020 (Corona et al. 2020). The ties were broken using this formula:

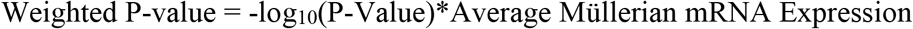

The same list was used for subsequent gene set enrichment analysis.

### Pathway gene set enrichment analysis

Pathway enrichment analysis was conducted using the ClusterProfiler package in R. We removed the HLA genes defined by the HLA Informatics Group (Tiercy et al. 2002; Hollenbach et al. 2011; Nunes et al. 2011) from the ranked list prior to carrying out gene set enrichment analysis. This is because the strong, long-distance linkage disequilibrium between SNPs in this region led to a clustering of multiple gene-level associations in this region making it difficult to differentiate between these genes in terms of ranks. This clustering in turn may yield potentially spurious enrichment signals for pathways that contain several HLA genes. We ran this analysis using the following script:

~~~
gseGO(geneList= <GENE-LIST>,
      ont =“ALL”,
      keyType = “ENTREZID”,
      nPerm = 10000,
      minGSSize = 3,
      maxGSSize = 800,
      pvalueCutoff = 0.05,
      verbose = TRUE,
      OrgDb = org.Hs.eg.db,
      pAdjustMethod = “BH”)
~~~

Pathways with gene set sizes less than 25 (as recommended by the BROAD institute) were removed from further analysis as the normalization to variation in gene set size becomes inaccurate for small gene sets.

### Super-enhancer associated TF gene set enrichment analysis

This analysis was conducted using the default GSEA preranked setting within the Broad GSEA (v3.0) program. The super-enhancer associated TF gene set was generated by taking known TFs from the Human Transcription Factor data base (Lambert et al. 2018), which was then filtered to only include TFs that were proximal, overlapping, or nearest to a super-enhancer as defined by ROSE2 (Whyte et al. 2013) for CCOC, EnOC, HGSOC, MOC, and NMOC (Corona et al. 2020). Enrichment plots were generated with the R package fgsea.

### MsigDB TF cistrome gene set enrichment analysis

This analysis was conducted using the default GSEA preranked setting within the Broad GSEA program (v3.0).

## EXPERIMENTAL METHODS

### Fallopian tube secretory epithelium RNA-seq

RNA-sequencing data from primary fallopian tube secretory epithelial cells were generated as described in Corona et al. 2020 (Corona et al. 2020). They are available in the GEO database under the accession code GSE182510.

## ACKNOWLEDGEMENTS

This project was supported by an Ovarian Cancer Research Fund Alliance Liz Tilberis Early Career Award (599175) (K.L.), Ovarian Cancer Research Fund Alliance Program Project Development (373356) (K.L.), a Southern California Clinical and Translational Science Institute Core Voucher (V148) (K.L.). SK is supported by a United Kingdom Research and Innovation Future Leaders Fellowship (MR/T043202/1). The research described was supported in part by NIH/National Center for Advancing Translational Science (NCATS) UCLA CTSI Grant Number UL1TR001881. R.N. is supported in part by a Ruth L. Kirschstein Institutional National Research Service Award (T32) from the NIH (grant number 5 T32 GM 118288-2). Funding details for the OCAC data set used in this manuscript are to be found in Coetzee, et al. 2021 (under review), ‘Integrative multi-omics analyses to identify the genetic and functional mechanisms underlying ovarian cancer risk regions’. NHS/NHS II was supported by NIH grants UM1 CA186107, P01 CA87969, R01 CA49449, U01 CA176726, and R01 CA67262. The content is solely the responsibility of the authors and does not necessarily represent the official views of the National Institutes of Health. NHS/NHS II thank the following state cancer registries for their help: AL, AZ, AR, CA, CO, CT, DE, FL, GA, ID, IL, IN, IA, KY, LA, ME, MD, MA, MI, NE, NH, NJ, NY, NC, ND, OH, OK, OR, PA, RI, SC, TN, TX, VA, WA, WY. The authors assume full responsibility for analyses and interpretation of these data. The NHS/NHS II study protocol was approved by the institutional review boards of the Brigham and Women’s Hospital and Harvard T.H. Chan School of Public Health, and those of participating registries as required. The NHS/NHS II acknowledge the Channing Division of Network Medicine, Department of Medicine, Brigham and Women’s Hospital and Harvard Medical School, Boston, MA, USA as the home of the Nurses’ Health Studies. We also thank the following individuals and institutions from the Ovarian Cancer Association Consortium - Simon G. Coetzee, Pei-Chen Peng, Will Rosenow, Stephanie Chen, Brian D. Davis, Felipe Segato Dezem, Ji-Heui Seo, Robbin Nameki, Alberto L. Reyes, Katja K.H. Aben, Natalia N. Antonenkova, Gerasimos Aravantinos, Laura E. Beane Freeman, Matthias W. Beckmann, Alicia Beeghly-Fadiel, Marcus Q. Bernardini, Line Bjorge, Amanda Black, Natalia V. Bogdanova, Kelly L. Bolton, Agnieszka Budzilowska, Ralf Butzow, Hui Cai, Rikki Cannioto, Kexin Chen, AOCS Group, Yoke-Eng Chiew, Linda S. Cook, Anna DeFazio, Joe Dennis, Jennifer A. Doherty, Thilo Dörk, Andreas du Bois, Diana M. Eccles, Gabrielle Ene, Peter A. Fasching, James M. Flanagan, Florentia Fostira, Aleksandra Gentry-Maharaj, Marc T. Goodman, Christopher A. Haiman, Florian Heitz, Michelle A.T. Hildebrandt, Estrid Høgdall, Claus K. Høgdall, Ruea-Yea Huang, Michael E. Jones, Daehee Kang, Beth Y. Karlan, Anthony N. Karnezis, Linda E. Kelemen, Catherine J. Kennedy, Elza K. Khusnutdinova, Lambertus A. Kiemeney, Susanne K. Kjaer, Jolanta Kupryjanczyk, Diether Lambrechts, Melissa C. Larson, Nhu D. Le, Jenny Lester, Lian Li, Jan Lubiński, Michael Lush, Keitaro Matsuo, Taymaa May, John R. McLaughlin, Francesmary Modugno, Melissa Moffitt, Steven A. Narod, Tu Nguyen-Dumont, Håkan Olsson, N. Charlotte Onland-Moret, Sue K. Park, Jennifer B. Permuth, Darya Prokofyeva, Harvey A. Risch, Cristina Rodríguez-Antona, V. Wendy Setiawan, Kang Shan, Melissa C. Southey, Anthony J. Swerdlow, Soo Hwang Teo, Kathryn L. Terry, Pamela J. Thompson, Liv Cecilie Vestrheim Thomsen, Cecilie F. Torkildsen, Linda Titus, Britton Trabert, Ruth Travis, Shelley S. Tworoger, Els Van Nieuwenhuysen, Digna Velez Edwards, Robert A. Vierkant, Rayna Matsuno Weise, Nicolas Wentzensen, Stacey J. Winham, Yin-Ling Woo, Li Yan, Wei Zheng, Argyrios Ziogas, Andrew Berchuck, Ellen L. Goode, Celeste L. Pearce, Susan J. Ramus, Thomas A. Sellers, Matthew L. Freedman, Joellen M. Schildkraut, Simon A. Gayther, Dennis Hazelett, Michelle R. Jones, Jasmine T. Plummer, MCCS, WMH. We also thank Pak Hin Yu for his technical assistance. Gene-essentiality results are generated by DepMap, BROAD https://doi.org/10.6084/m9.figshare.12280541.v4.

## AUTHOR CONTRIBUTIONS

Directed the study R.N., A.S., K.L., S.K. Lead data analyst, data curation: R.N., A.S., R.C. Data generation, data analyst: E.D., J.T., P.P., OCAC., X.L. Conceived, designed, and supervised the project: K.L., S.K. Drafted the manuscript: R.N., A.S., K.L., S.K. All authors reviewed and approved the final draft.

## ACCESSION CODES

GSE182510

## CODE AVAILABILITY

A step-by-step tutorial of chromMAGMA is available in a Github repository: https://github.com/lawrenson-lab/chromMAGMA-public.

## BIBLIOGRAPHY

1000 Genomes Project Consortium, Auton A, Brooks LD, Durbin RM, Garrison EP, Kang HM, Korbel JO, Marchini JL, McCarthy S, McVean GA, et al. 2015. A global reference for human genetic variation. Nature 526: 68–74.

Adler EK, Corona RI, Lee JM, Rodriguez-Malave N, Mhawech-Fauceglia P, Sowter H, Hazelett DJ, Lawrenson K, Gayther SA. 2017. The PAX8 cistrome in epithelial ovarian cancer. Oncotarget 8: 108316–108332.

Berisa T, Pickrell JK. 2016. Approximately independent linkage disequilibrium blocks in human populations. Bioinformatics 32: 283–285.

Bleu M, Mermet-Meillon F, Apfel V, Barys L, Holzer L, Bachmann Salvy M, Lopes R, Amorim Monteiro Barbosa I, Delmas C, Hinniger A, et al. 2021. PAX8 and MECOM are interaction partners driving ovarian cancer. Nat Commun 12: 2442.

Bojesen SE, Pooley KA, Johnatty SE, Beesley J, Michailidou K, Tyrer JP, Edwards SL, Pickett HA, Shen HC, Smart CE, et al. 2013. Multiple independent variants at the TERT locus are associated with telomere length and risks of breast and ovarian cancer. Nat Genet 45: 371–84, 384e1.

Bolton KL, Tyrer J, Song H, Ramus SJ, Notaridou M, Jones C, Sher T, Gentry-Maharaj A, Wozniak E, Tsai Y-Y, et al. 2010. Common variants at 19p13 are associated with susceptibility to ovarian cancer. Nat Genet 42: 880–884.

Boyle EA, Li YI, Pritchard JK. 2017. An expanded view of complex traits: from polygenic to omnigenic. Cell 169: 1177–1186.

Bradner JE, Hnisz D, Young RA. 2017. Transcriptional addiction in cancer. Cell 168: 629–643.

Coetzee S, Dareng EO, Peng P, Rosenow W, Tyrer JP. 2021. Integrative multi-omics analyses to identify the genetic and functional mechanisms underlying ovarian cancer risk regions. (under review)

Coetzee SG, Shen HC, Hazelett DJ, Lawrenson K, Kuchenbaecker K, Tyrer J, Rhie SK, Levanon K, Karst A, Drapkin R, et al. 2015. Cell-type-specific enrichment of risk-associated regulatory elements at ovarian cancer susceptibility loci. Hum Mol Genet 24: 3595–3607.

Corces MR, Granja JM, Shams S, Louie BH, Seoane JA, Zhou W, Silva TC, Groeneveld C, Wong CK, Cho SW, et al. 2018. The chromatin accessibility landscape of primary human cancers. Science 362.

Corona RI, Seo J-H, Lin X, Hazelett DJ, Reddy J, Abassi F, Lin YG, Mhawech-Fauceglia PY, Lester J, Shah SP, et al. 2019. Non-coding Somatic Mutations Converge on the PAX8 Pathway in Epithelial Ovarian Cancer. BioRxiv.

Corona RI, Seo J-H, Lin X, Hazelett DJ, Reddy J, Fonseca MAS, Abassi F, Lin YG, Mhawech-Fauceglia PY, Shah SP, et al. 2020. Non-coding somatic mutations converge on the PAX8 pathway in ovarian cancer. Nat Commun 11: 2020.

Cuff J, Salari K, Clarke N, Esheba GE, Forster AD, Huang S, West RB, Higgins JP, Longacre TA, Pollack JR. 2013. Integrative bioinformatics links HNF1B with clear cell carcinoma and tumor-associated thrombosis. PLoS ONE 8: e74562.

de Leeuw CA, Mooij JM, Heskes T, Posthuma D. 2015. MAGMA: generalized gene-set analysis of GWAS data. PLoS Comput Biol 11: e1004219.

Demontis D, Walters RK, Martin J, Mattheisen M, Als TD, Agerbo E, Baldursson G, Belliveau R, Bybjerg-Grauholm J, Bækvad-Hansen M, et al. 2019. Discovery of the first genome-wide significant risk loci for attention deficit/hyperactivity disorder. Nat Genet 51: 63–75.

Fishilevich S, Nudel R, Rappaport N, Hadar R, Plaschkes I, Iny Stein T, Rosen N, Kohn A, Twik M, Safran M, et al. 2017. GeneHancer: genome-wide integration of enhancers and target genes in GeneCards. Database (Oxford) 2017.

Fulco CP, Nasser J, Jones TR, Munson G, Bergman DT, Subramanian V, Grossman SR, Anyoha R, Doughty BR, Patwardhan TA, et al. 2019. Activity-by-contact model of enhancer-promoter regulation from thousands of CRISPR perturbations. Nat Genet 51: 1664–1669.

Garcia-Alonso L, Handfield L-F, Roberts K, Nikolakopoulou K, Fernando RC, Gardner L, Woodhams B, Arutyunyan A, Polanski K, Hoo R, et al. 2021. Mapping the temporal and spatial dynamics of the human endometrium in vivo and in vitro. Nat Genet 53: 1698–1711.

Gerasimova A, Chavez L, Li B, Seumois G, Greenbaum J, Rao A, Vijayanand P, Peters B. 2013. Predicting cell types and genetic variations contributing to disease by combining GWAS and epigenetic data. PLoS ONE 8: e54359.

Goode EL, Chenevix-Trench G, Song H, Ramus SJ, Notaridou M, Lawrenson K, Widschwendter M, Vierkant RA, Larson MC, Kjaer SK, et al. 2010. A genome-wide association study identifies susceptibility loci for ovarian cancer at 2q31 and 8q24. Nat Genet 42: 874–879.

Grisanzio C, Freedman ML. 2010. Chromosome 8q24-Associated Cancers and MYC. Genes Cancer 1: 555–559.

Gusev A, Lawrenson K, Lin X, Lyra PC, Kar S, Vavra KC, Segato F, Fonseca MAS, Lee JM, Pejovic T, et al. 2019. A transcriptome-wide association study of high-grade serous epithelial ovarian cancer identifies new susceptibility genes and splice variants. Nat Genet 51: 815–823.

Hay D, Hughes JR, Babbs C, Davies JOJ, Graham BJ, Hanssen L, Kassouf MT, Marieke Oudelaar AM, Sharpe JA, Suciu MC, et al. 2016. Genetic dissection of the α-globin super-enhancer in vivo. Nat Genet 48: 895–903.

Hnisz D, Abraham BJ, Lee TI, Lau A, Saint-André V. 2013. Transcriptional super-enhancers connected to cell identity and disease. Cell.

Hollenbach JA, Mack SJ, Gourraud PA, Single RM, Maiers M, Middleton D, Thomson G, Marsh SGE, Varney MD, Immunogenomics Data Analysis Working Group. 2011. A community standard for immunogenomic data reporting and analysis: proposal for a STrengthening the REporting of Immunogenomic Studies statement. Tissue Antigens 78: 333–344.

Huang J, Li K, Cai W, Liu X, Zhang Y, Orkin SH, Xu J, Yuan G-C. 2018. Dissecting super-enhancer hierarchy based on chromatin interactions. Nat Commun 9: 943.

Huo Y, Li S, Liu J, Li X, Luo X-J. 2019. Functional genomics reveal gene regulatory mechanisms underlying schizophrenia risk. Nat Commun 10: 670.

Idaikkadar P, Morgan R, Michael A. 2019. HOX genes in high grade ovarian cancer. Cancers (Basel) 11.

Jansen IE, Savage JE, Watanabe K, Bryois J, Williams DM, Steinberg S, Sealock J, Karlsson IK, Hägg S, Athanasiu L, et al. 2019. Genome-wide meta-analysis identifies new loci and functional pathways influencing Alzheimer’s disease risk. Nat Genet 51: 404–413.

Jones MR, Kamara D, Karlan BY, Pharoah PDP, Gayther SA. 2017. Genetic epidemiology of ovarian cancer and prospects for polygenic risk prediction. Gynecol Oncol 147: 705–713.

Jones MR, Peng P-C, Coetzee SG, Tyrer J, Reyes ALP, Corona RI, Davis B, Chen S, Dezem F, Seo J-H, et al. 2020. Ovarian Cancer Risk Variants Are Enriched in Histotype-Specific Enhancers and Disrupt Transcription Factor Binding Sites. Am J Hum Genet 107: 622–635.

Kandaswamy R, Sava GP, Speedy HE, Beà S, Martín-Subero JI, Studd JB, Migliorini G, Law PJ, Puente XS, Martín-García D, et al. 2016. Genetic Predisposition to Chronic Lymphocytic Leukemia Is Mediated by a BMF Super-Enhancer Polymorphism. Cell Rep 16: 2061–2067.

Kar S, Considine D, Tyrer J, Plummer J, Chen S, Dezem F, Barbeira A, Rajagopal P, Rosenow W, Anton F, et al. 2020. Pleiotropy-guided transcriptome imputation from normal and tumor tissues identifies new candidate susceptibility genes for breast and ovarian cancer. BioRxiv.

Kar SP, Adler E, Tyrer J, Hazelett D, Anton-Culver H, Bandera EV, Beckmann MW, Berchuck A, Bogdanova N, Brinton L, et al. 2017. Enrichment of putative PAX8 target genes at serous epithelial ovarian cancer susceptibility loci. Br J Cancer 116: 524–535.

Kar SP, Beesley J, Amin Al Olama A, Michailidou K, Tyrer J, Kote-Jarai Zs, Lawrenson K, Lindstrom S, Ramus SJ, Thompson DJ, et al. 2016. Genome-Wide Meta-Analyses of Breast, Ovarian, and Prostate Cancer Association Studies Identify Multiple New Susceptibility Loci Shared by at Least Two Cancer Types. Cancer Discov 6: 1052–1067.

Kawakami R, Kitagawa Y, Chen KY, Arai M, Ohara D, Nakamura Y, Yasuda K, Osaki M, Mikami N, Lareau CA, et al. 2021. Distinct Foxp3 enhancer elements coordinate development, maintenance, and function of regulatory T cells. Immunity 54: 947–961.e8.

Kelemen LE, Lawrenson K, Tyrer J, Li Q, Lee JM, Seo J-H, Phelan CM, Beesley J, Chen X, Spindler TJ, et al. 2015. Genome-wide significant risk associations for mucinous ovarian carcinoma. Nat Genet 47: 888–897.

Kim W, Bird GH, Neff T, Guo G, Kerenyi MA, Walensky LD, Orkin SH. 2013. Targeted disruption of the EZH2-EED complex inhibits EZH2-dependent cancer. Nat Chem Biol 9: 643–650.

Kuchenbaecker KB, Ramus SJ, Tyrer J, Lee A, Shen HC, Beesley J, Lawrenson K, McGuffog L, Healey S, Lee JM, et al. 2015. Identification of six new susceptibility loci for invasive epithelial ovarian cancer. Nat Genet 47: 164–171.

Lambert SA, Jolma A, Campitelli LF, Das PK, Yin Y, Albu M, Chen X, Taipale J, Hughes TR, Weirauch MT. 2018. The human transcription factors. Cell 172: 650–665.

Lawrenson K, Kar S, McCue K, Kuchenbaeker K, Michailidou K, Tyrer J, Beesley J, Ramus SJ, Li Q, Delgado MK, et al. 2016. Functional mechanisms underlying pleiotropic risk alleles at the 19p13.1 breast-ovarian cancer susceptibility locus. Nat Commun 7: 12675.

Lawrenson K, Li Q, Kar S, Seo J-H, Tyrer J, Spindler TJ, Lee J, Chen Y, Karst A, Drapkin R, et al. 2015. Cis-eQTL analysis and functional validation of candidate susceptibility genes for high-grade serous ovarian cancer. Nat Commun 6: 8234.

Li Q, Zeng X, Cheng X, Zhang J, Ji J, Wang J, Xiong K, Qi Q, Huang W. 2015. Diagnostic value of dual detection of hepatocyte nuclear factor 1 beta (HNF-1β) and napsin A for diagnosing ovarian clear cell carcinoma. Int J Clin Exp Pathol 8: 8305–8310.

Liberzon A, Subramanian A, Pinchback R, Thorvaldsdóttir H, Tamayo P, Mesirov JP. 2011. Molecular signatures database (MsigDB) 3.0. Bioinformatics 27: 1739–1740.

Lu Y, Beeghly-Fadiel A, Wu L, Guo X, Li B, Schildkraut JM, Im HK, Chen YA, Permuth JB, Reid BM, et al. 2018. A Transcriptome-Wide Association Study Among 97,898 Women to Identify Candidate Susceptibility Genes for Epithelial Ovarian Cancer Risk. Cancer Res 78: 5419–5430.

Manolio TA, Collins FS, Cox NJ, Goldstein DB, Hindorff LA, Hunter DJ, McCarthy MI, Ramos EM, Cardon LR, Chakravarti A, et al. 2009. Finding the missing heritability of complex diseases. Nature 461: 747–753.

Meyers RM, Bryan JG, McFarland JM, Weir BA, Sizemore AE, Xu H, Dharia NV, Montgomery PG, Cowley GS, Pantel S, et al. 2017. Computational correction of copy number effect improves specificity of CRISPR-Cas9 essentiality screens in cancer cells. Nat Genet 49: 1779–1784.

Muerdter F, Boryń ŁM, Arnold CD. 2015. STARR-seq-principles and applications. Genomics 106: 145–150.

Nameki R, Chang H, Reddy J, Corona RI, Lawrenson K. 2021. Transcription factors in epithelial ovarian cancer: histotype-specific drivers and novel therapeutic targets. Pharmacol Ther 220: 107722.

Nasser J, Bergman DT, Fulco CP, Guckelberger P, Doughty BR, Patwardhan TA, Jones TR, Nguyen TH, Ulirsch JC, Lekschas F, et al. 2021. Genome-wide enhancer maps link risk variants to disease genes. Nature 593: 238–243.

Nunes E, Heslop H, Fernandez-Vina M, Taves C, Wagenknecht DR, Eisenbrey AB, Fischer G, Poulton K, Wacker K, Hurley CK, et al. 2011. Definitions of histocompatibility typing terms. Blood 118: e180–3.

Oldridge DA, Wood AC, Weichert-Leahey N, Crimmins I, Sussman R, Winter C, McDaniel LD, Diamond M, Hart LS, Zhu S, et al. 2015. Genetic predisposition to neuroblastoma mediated by a LMO1 super-enhancer polymorphism. Nature 528: 418–421.

Peng Y, Zhang Y. 2018. Enhancer and super-enhancer: Positive regulators in gene transcription. Anim Models Exp Med 1: 169–179.

Pharoah PDP, Tsai Y-Y, Ramus SJ, Phelan CM, Goode EL, Lawrenson K, Buckley M, Fridley BL, Tyrer JP, Shen H, et al. 2013. GWAS meta-analysis and replication identifies three new susceptibility loci for ovarian cancer. Nat Genet 45: 362–70, 370e1.

Phelan CM, Kuchenbaecker KB, Tyrer JP, Kar SP, Lawrenson K, Winham SJ, Dennis J, Pirie A, Riggan MJ, Chornokur G, et al. 2017. Identification of 12 new susceptibility loci for different histotypes of epithelial ovarian cancer. Nat Genet 49: 680–691.

Quinlan AR, Hall IM. 2010. BEDTools: a flexible suite of utilities for comparing genomic features. Bioinformatics 26: 841–842.

Reddy J, Fonseca MAS, Corona RI, Nameki R, Segato Dezem F, Klein IA, Chang H, Chaves-Moreira D, Afeyan L, Malta TM, et al. 2019. Predicting master transcription factors from pan-cancer expression data. BioRxiv.

Sanyal A, Lajoie BR, Jain G, Dekker J. 2012. The long-range interaction landscape of gene promoters. Nature 489: 109–113.

Sey NYA, Hu B, Mah W, Fauni H, McAfee JC, Rajarajan P, Brennand KJ, Akbarian S, Won H. 2020. A computational tool (H-MAGMA) for improved prediction of brain-disorder risk genes by incorporating brain chromatin interaction profiles. Nat Neurosci 23: 583–593.

Shin HY, Willi M, HyunYoo K, Zeng X, Wang C, Metser G, Hennighausen L. 2016. Hierarchy within the mammary STAT5-driven Wap super-enhancer. Nat Genet 48: 904–911.

Smedley D, Haider S, Durinck S, Pandini L, Provero P, Allen J, Arnaiz O, Awedh MH, Baldock R, Barbiera G, et al. 2015. The BioMart community portal: an innovative alternative to large, centralized data repositories. Nucleic Acids Res 43: W589–98.

Song H, Ramus SJ, Tyrer J, Bolton KL, Gentry-Maharaj A, Wozniak E, Anton-Culver H, Chang-Claude J, Cramer DW, DiCioccio R, et al. 2009. A genome-wide association study identifies a new ovarian cancer susceptibility locus on 9p22.2. Nat Genet 41: 996–1000.

Soslow RA. 2008. Histologic subtypes of ovarian carcinoma: an overview. Int J Gynecol Pathol 27: 161–174.

Subramanian A, Tamayo P, Mootha VK, Mukherjee S, Ebert BL, Gillette MA, Paulovich A, Pomeroy SL, Golub TR, Lander ES, et al. 2005. Gene set enrichment analysis: a knowledge-based approach for interpreting genome-wide expression profiles. Proc Natl Acad Sci USA 102:15545–15550.

Tiercy JM, Marsh SGE, Schreuder GMT, Albert E, Fischer G, Wassmuth R, European Federation for Immunogenetics subcommittee for reporting HLA ambiguities. 2002. Guidelines for nomenclature usage in HLA reports: ambiguities and conversion to serotypes. Eur J Immunogenet 29: 273–274.

Torre LA, Trabert B, DeSantis CE, Miller KD, Samimi G, Runowicz CD, Gaudet MM, Jemal A, Siegel RL. 2018. Ovarian cancer statistics, 2018. CA Cancer J Clin 68: 284–296.

Vizcaíno C, Mansilla S, Portugal J. 2015. Sp1 transcription factor: A long-standing target in cancer chemotherapy. Pharmacol Ther 152: 111–124.

Whyte WA, Orlando DA, Hnisz D, Abraham BJ, Lin CY, Kagey MH, Rahl PB, Lee TI, Young RA. 2013. Master transcription factors and mediator establish super-enhancers at key cell identity genes. Cell 153: 307–319.

Wray NR, Ripke S, Mattheisen M, Trzaskowski M, Byrne EM, Abdellaoui A, Adams MJ, Agerbo E, Air TM, Andlauer TMF, et al. 2018. Genome-wide association analyses identify 44 risk variants and refine the genetic architecture of major depression. Nat Genet 50: 668–681.

Zhang F, Lupski JR. 2015. Non-coding genetic variants in human disease. Hum Mol Genet 24: R102–10.

2019. 12 Impact of functional information on understanding variation. Nature.

